## Supplementary files for "Umbilical cord blood-derived cell therapy for perinatal brain injury: A systematic review & meta-analysis of preclinical studies - Part A"

**Table S1 – Search strategy & search terms**

| **#** | **Query** |
| --- | --- |
| 1 | (perinatal or neonat* or newborn or infant or f?et* or prematur*).tw. |
| 2 | exp Animals/ or exp Rats/ or exp Mice/ or exp Sheep/ or exp Swine/ or exp Rabbits/ or exp Disease Models, Animal/ |
| 3 | 1 and 2 |
| 4 | (brain or cerebral or encephalopath* or neuro*).tw. |
| 5 | (injury or damage or insult or trauma or hypox* or anox* or oxygen deprivation or isch?em* or h?emorrhag* or stroke or thrombosis or hypoperfusion or underperfusion or asphyxia or growth restriction or periventricular leukomalacia or inflamm* or chorioamnionitis or infection or sepsis or toxin or hypoglyc?em* or placental insufficiency or placental abnormality or placental lesion or placental abruption or uterine rupture or maternal obesity or metabolic or stress or distress or vascular dysfunction or low birth weight or cord prolapse or eclampsia or preeclampsia).tw. |
| 6 | 4 and 5 |
| 7 | (umbilical cord and cell).tw. |
| 8 | umbilical cord blood.tw. |
| 9 | (cell and cord blood).tw. |
| 10 | (h?ematopoietic and cord blood).tw. |
| 11 | (mesenchymal and cord blood).tw |
| 12 | (stromal and cord blood).tw. |
| 13 | (endothelial progenitor and cord blood).tw. |
| 14 | (mononuclear and cord blood).tw. |
| 15 | (regulatory T and cord blood).tw. |
| 16 | 7 or 8 or 9 or 10 or 11 or 12 or 13 or 14 or 15 |
| 17 | 3 and 6 and 16 |

**Figures S1 – Publication bias as assessed by funnel plot analysis & Egger’s test (A)** Infarct size; **(B)** Neuron number; **(C)** Oligodendrocyte number – grey matter**; (D)** Oligodendrocyte number – white matter; **(E)** Apoptosis – grey matter; **(F)** Apoptosis – white matter; **(G)** Astrogliosis – grey matter; **(H)** Astrogliosis – white matter; **(I)** Microglia activation – grey matter; **(J)** Microglia activation – white matter; **(K)** TNF-α; **(L)** IL-6; **(M)** IL-1 β; **(N)** IL-10; **(O)** Cylinder test; **(P)** Rotarod test

**(A)**

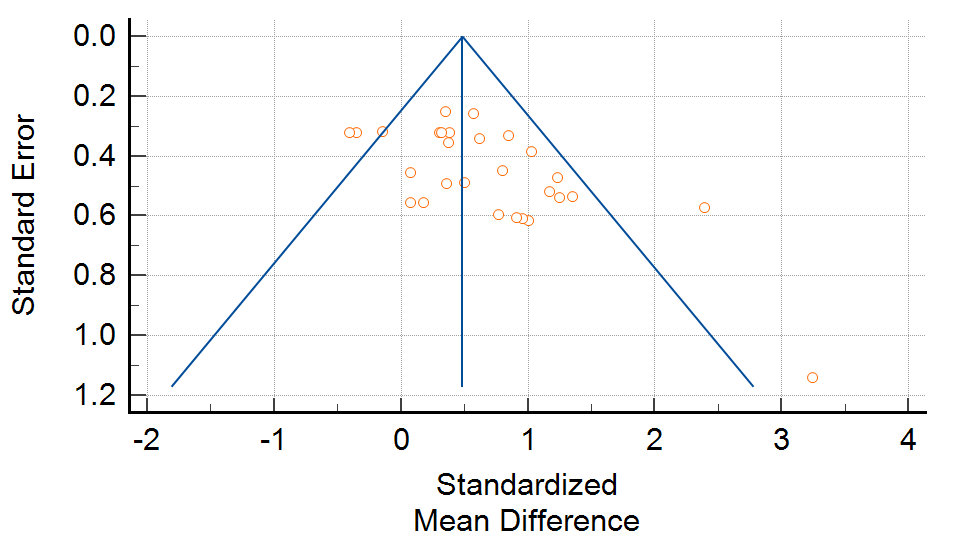

**Egger’s test:** 2.58 (1.04, 4.13), P = 0.0020

**(B)**

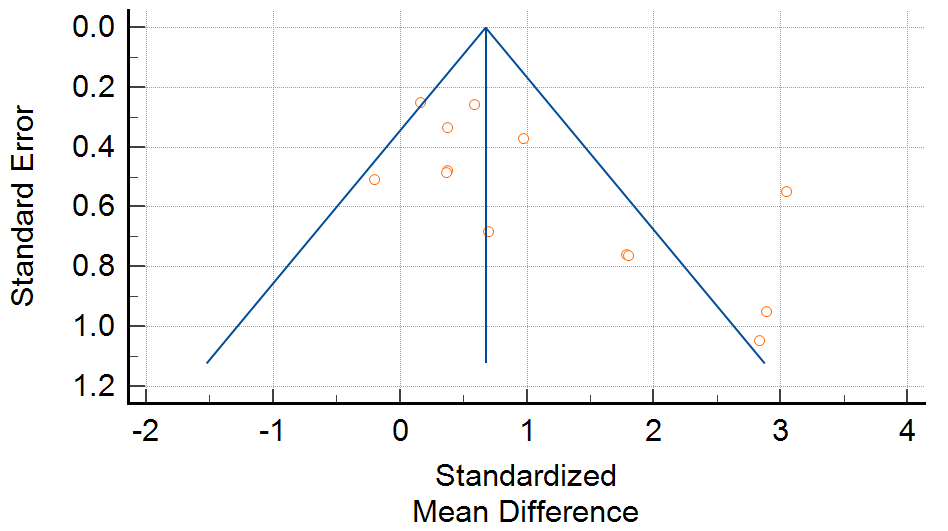

**Egger’s test:** 2.78 (0.54, 5.02), P = 0.019

**(C)**

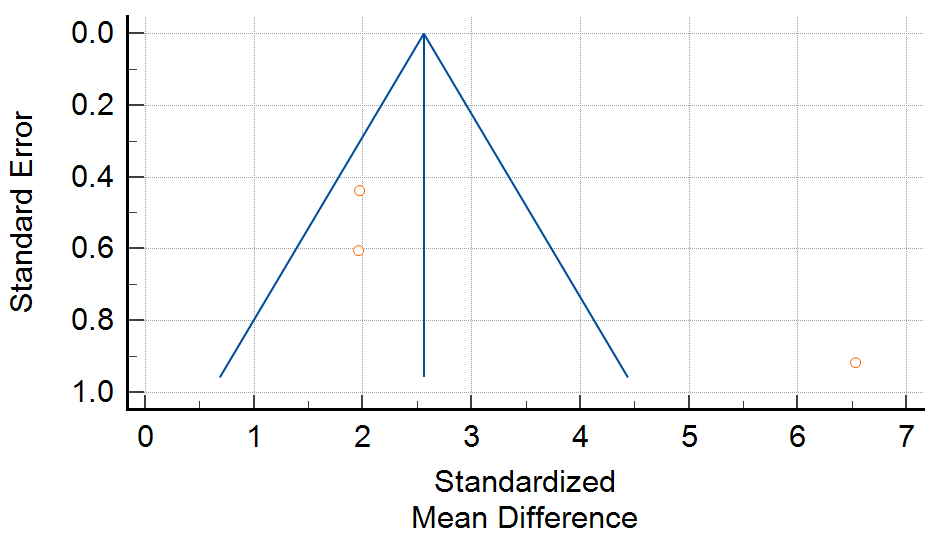

**Egger’s test:** 8.51 (-48.59, 65.51), P = 0.3089

**(D)**

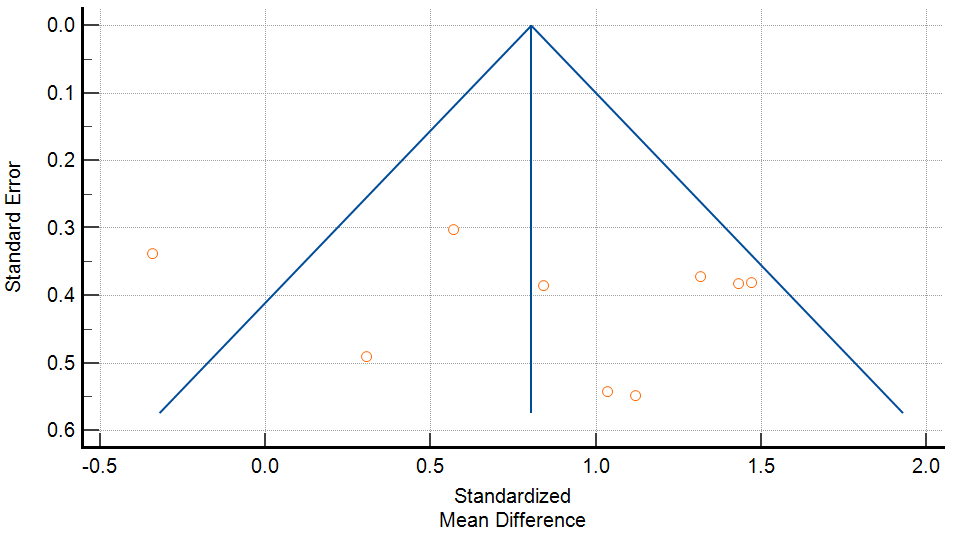

**Egger’s test:** 2.18 (-4.93, 9.29), P = 0.49

**(E)**

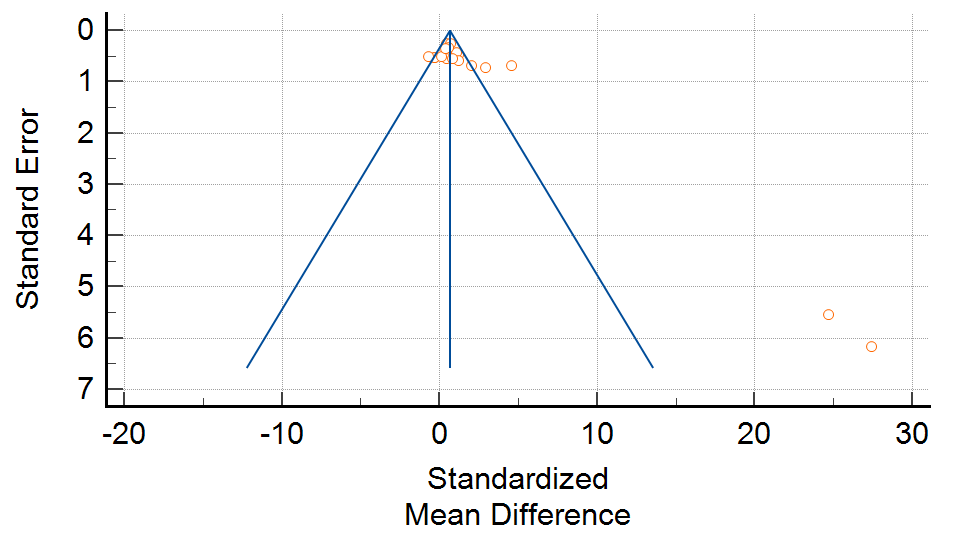

**Egger’s test:** 3.51 (1.37, 5.65), P = 0.0030

**(F)**

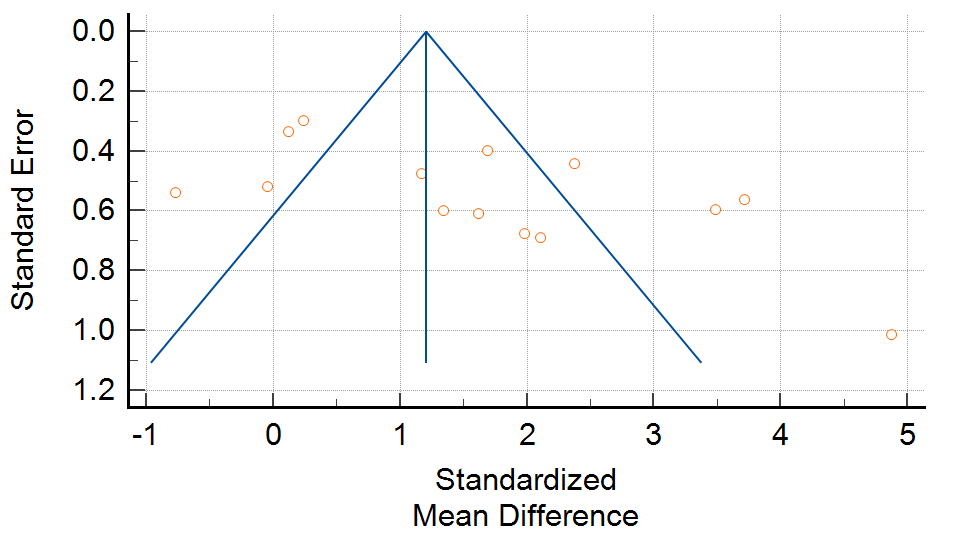

**Egger’s test:** 5.44 (0.83, 10.05), P = 0.025

**(G)**

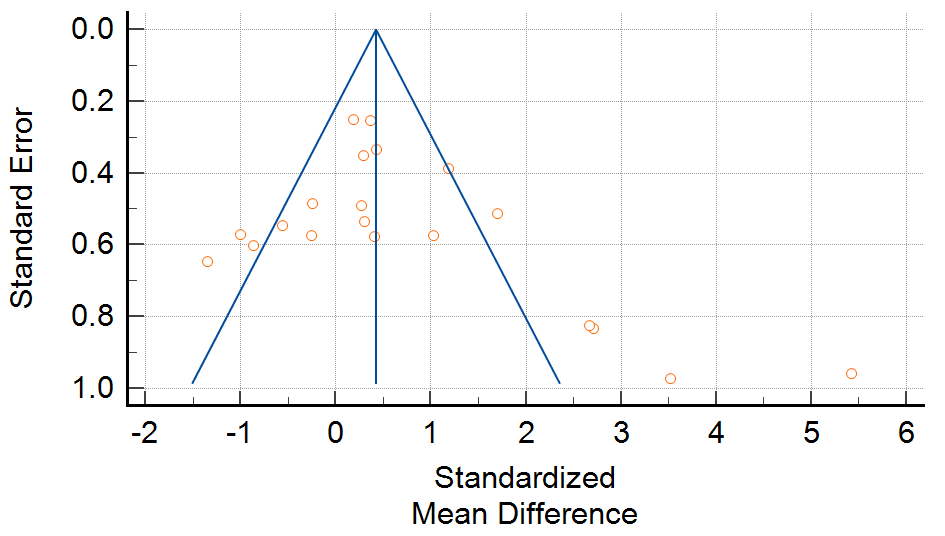

**Egger’s test:** 1.99 (-0.63, 4.61), P = 0.13

**(H)**

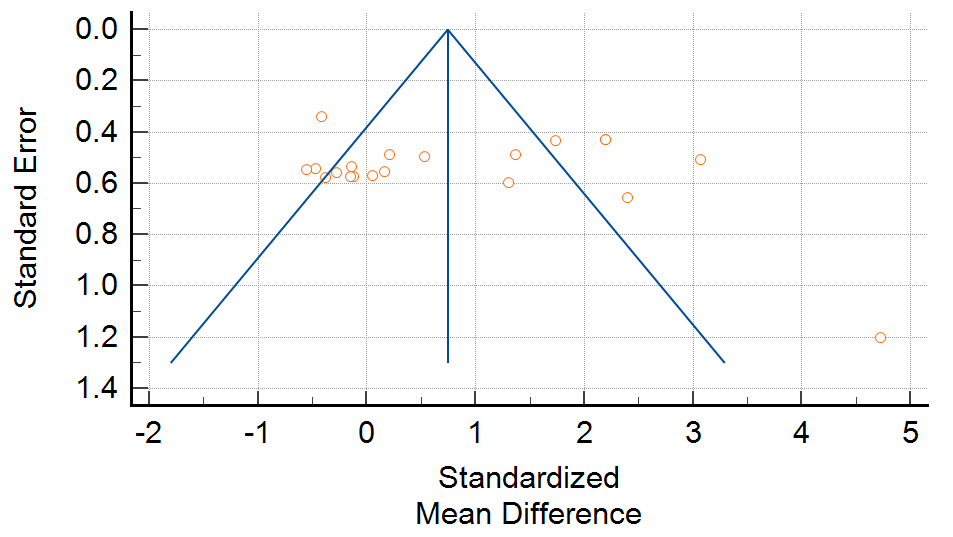

**Egger’s test:** 1.64 (-4.013, 7.30), P = 0.55

**(I)**

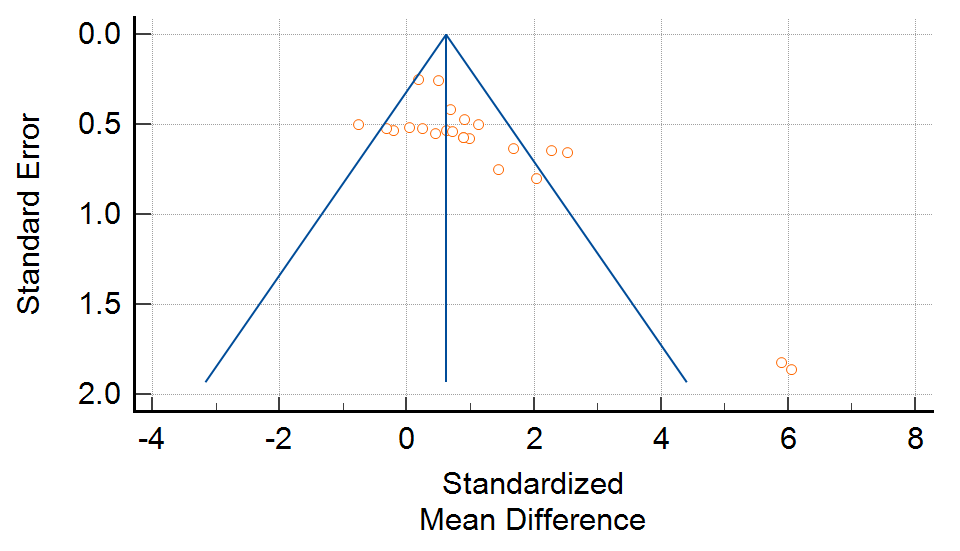

**Egger’s test:** 2.73 (1.23, 4.22), P = 0.0010

**(J)**

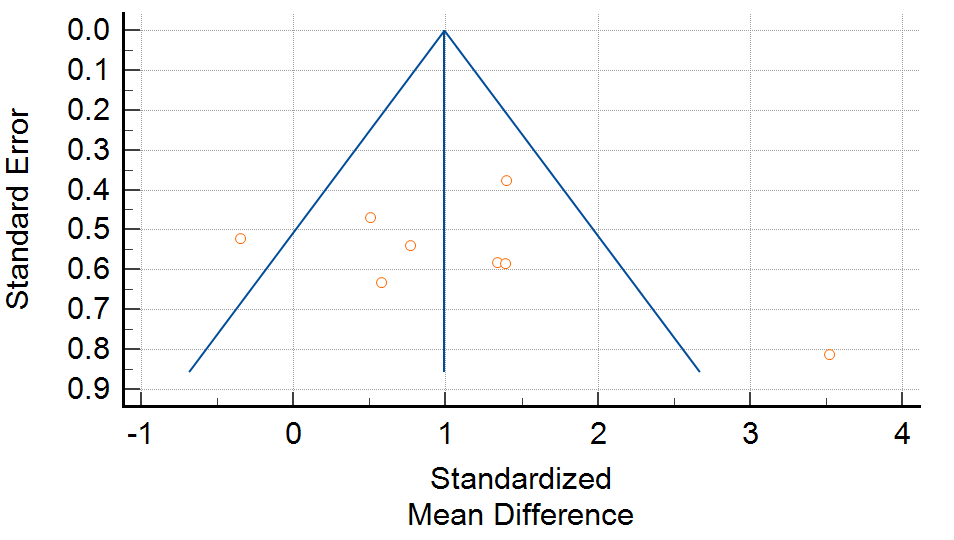

**Egger’s test:** 2.38 (-4.85, 9.62), P = 0.45

**(K)**

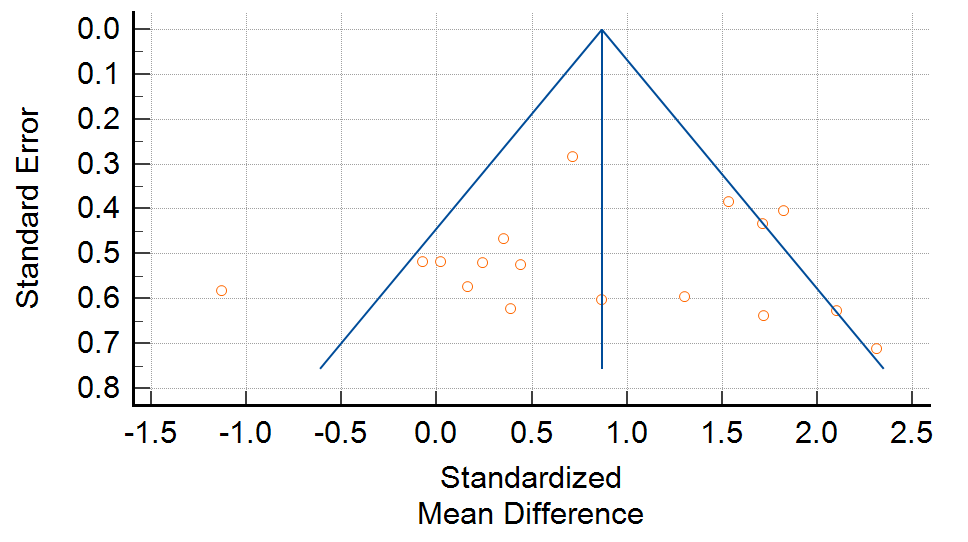

**Egger’s test:** -0.45 (-4.13, 3.22), P = 0.80

**(L)**

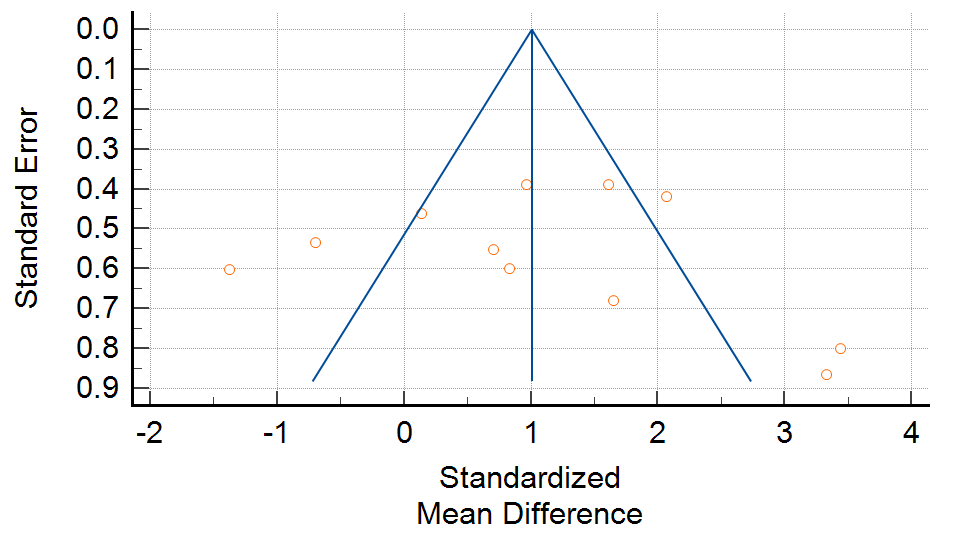

**Egger’s test:** 1.21 (-5.65, 8.08), P = 0.70

**(M)**

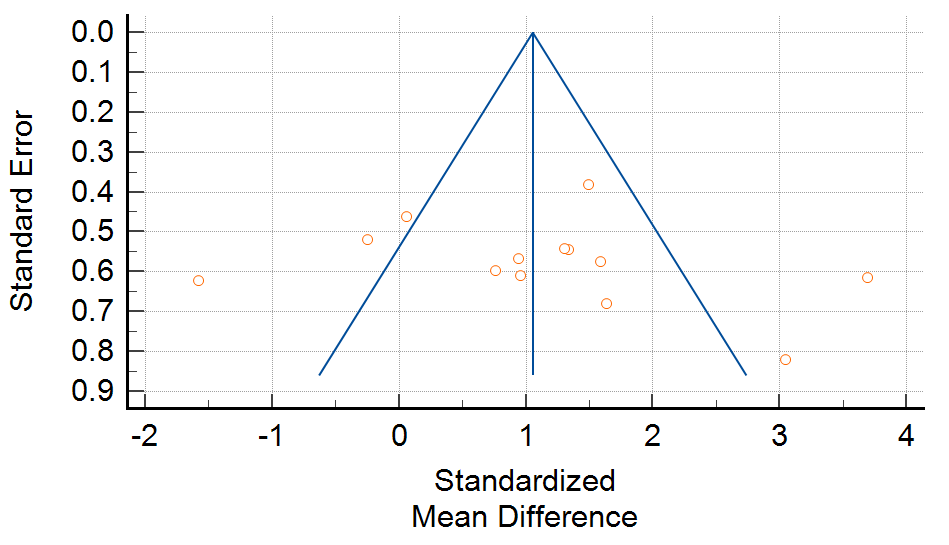

**Egger’s test:** 2.22 (-5.35, 9.79), P = 0.53

**(N)**

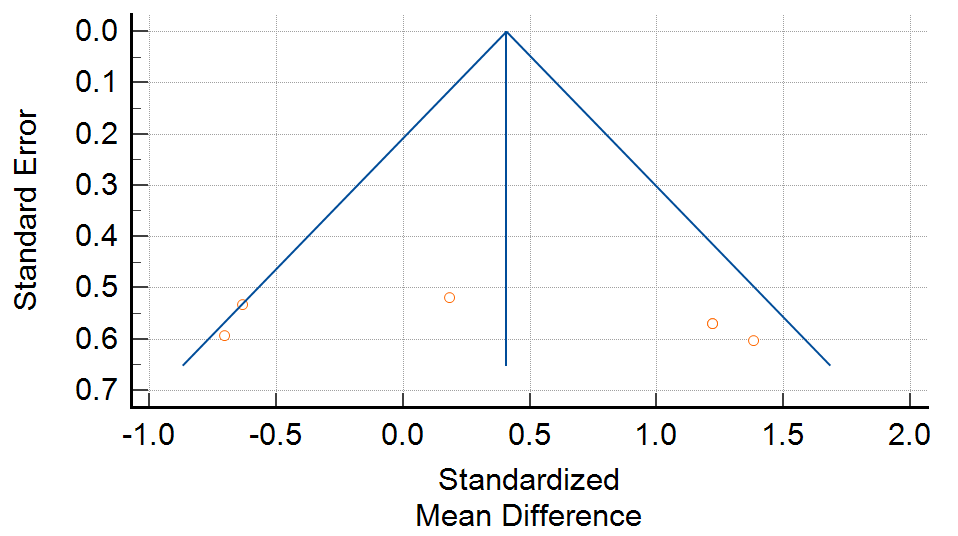

**Egger’s test:** 11.27 (-24.82, 47.35), P = 0.43

**(O)**

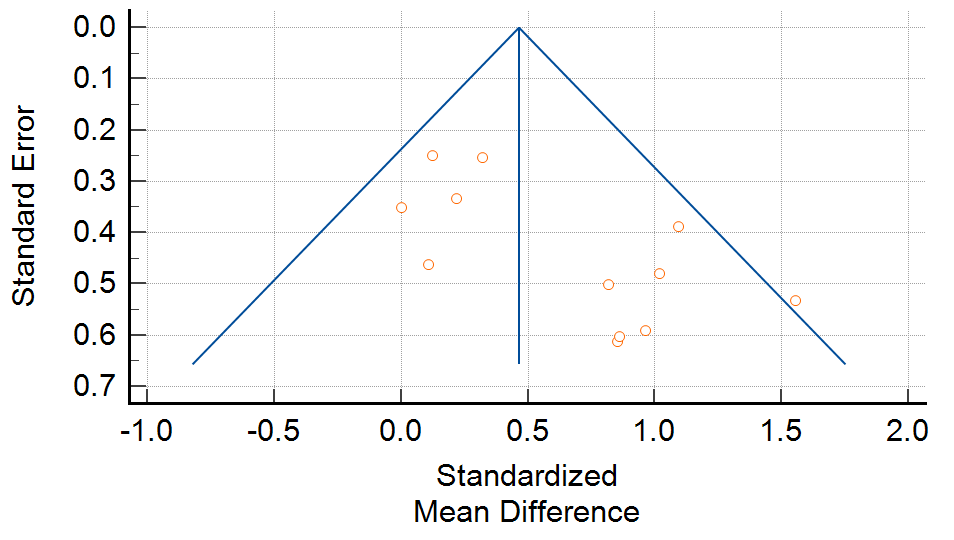

**Egger’s test:** 2.49 (0.64, 4.34), P = 0.013

**(P)**

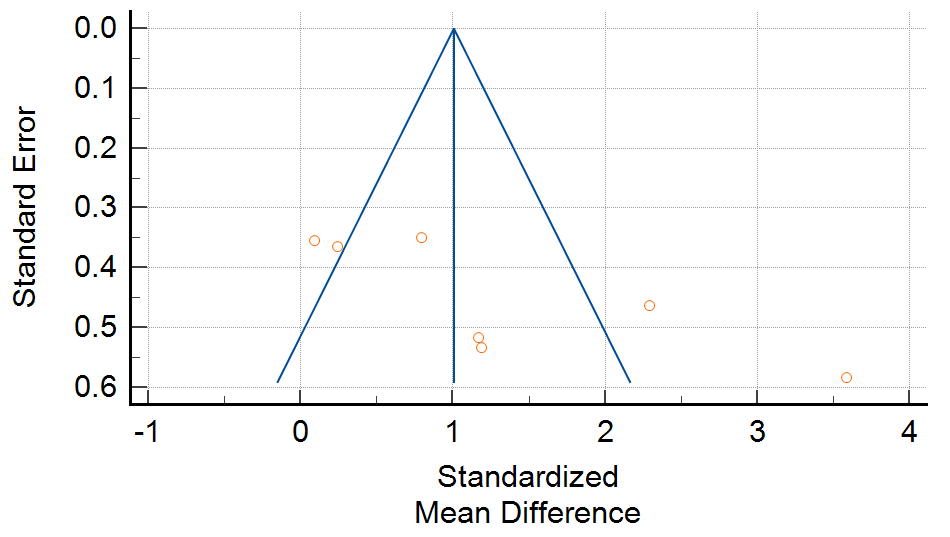

**Egger’s test:** 9.61 (1.12, 18.10), P = 0.033
